# A cost-effective and efficient approach for generating and assembling reagents for conducting real-time PCR

**DOI:** 10.1101/2021.07.14.452300

**Authors:** Ridim D Mote, Shinde Laxmikant V, Surya Bansi Singh, Mahak Tiwari, Hemant, Juhi Srivastava, Vidisha Tripathi, Vasudevan Seshadri, Amitabha Majumdar, Deepa Subramanyam

**Affiliations:** National Centre for Cell Science, SP Pune University Campus, Ganeshkhind, Pune – 411007, India; Savitribai Phule Pune University, Ganeshkhind, Pune – 411007, India; Dr. Babasaheb Ambedkar Marathwada University, Aurangabad – 431004, India; Applied Parasitology Research Laboratory, Department of Zoology, JES College, Jalna - 431203, India

**Author notes:** Corresponding authors: Deepa Subramanyam, Amitabha Majumdar.

**Keywords:** Real-time PCR, SYBR Green I, EvaGreen, Moloney Murine Leukemia Virus Reverse Transcriptase (M-MLV RT), hot-start Taq polymerase, in-house PCR mastermix

## Abstract

Real-time PCR is a widely used technique for quantification of gene expression. However, commercially available kits for real-time PCR are very expensive. The ongoing coronavirus pandemic has severely hampered the economy in a number of developing countries, resulting in a reduction in available research funding. The fallout of this will result in limiting educational institutes and small enterprises from using cutting edge biological techniques such as real-time PCR. Here, we report a cost-effective approach for preparing and assembling cDNA synthesis and real-time PCR mastermixes with similar efficiencies as commercially available kits. Our results thus demonstrate an alternative to commercially available kits.

## Introduction

Real-time polymerase chain reaction (Real-time PCR) is a powerful technique to measure the level of gene expression. It is a quantitative PCR technique where data is collected simultaneously as the PCR amplification proceeds. It is an extremely sensitive technique with a large dynamic range and high sequence specificity (Wong and Medrano 2005), with ability to even detect a single copy of a specific transcript (Palmer *et al.* 2003). Real-time PCR is characterized by a Ct (Cycle threshold) value which indicates the cycle where the fluorescence intensity of the PCR product is greater than the background fluorescence (Heid *et al.* 1996). Detection of the amplicon in real-time PCR can be done in multiple ways. The amplicon can be detected by using hybridization probes such as Taqman probes (Holland *et al.* 1991), molecular beacons (Tyagi and Kramer 1996), Eclipse Probes (Lukhtanov *et al.* 2007), LUX PCR Primers (Vilcek *et al.* 2010) and Scorpions (Whitcombe *et al.* 1999). While these probes are target sequence-specific, they are not commonly used due to their high cost. For the detection of a large number of genes, fluorescent DNA binding dyes, which are not sequence-specific, and intercalate between double-stranded DNA can be employed. Various DNA binding dyes such as SYBR Green I (Green and Sambrook 2018) and SYTO dyes (Gudnason *et al.* 2007; Eischeid 2011) are commercially used to detect the amplified PCR product, of which SYBR Green I is the most widely used. However, the commercially available SYBR Green I mastermixes are expensive. Hence, we set out with a goal to design a low-cost, in-house, real-time PCR mastermix using reagents easily available in labs, and which are as efficient as commercially available mastermixes.

SYBR Green I is the most widely used dye in real-time PCR mastermixes. Recently, EvaGreen dye has also been used for quantification in real-time PCR(Dhami and Kumarasinghe 2014). Reaction efficiency for EvaGreen has been shown to outperform SYBR Green I (Eischeid 2011). EvaGreen dye is spectrally similar to other dyes such as SYBR Green I and FAM (Mao *et al.* 2007). Hence, no changes are required in the optical settings of the instrument while using EvaGreen dye. Here, we demonstrate a protocol to purify in-house Moloney Murine Leukemia Virus Reverse Transcriptase (M-MLV RT) (Graham *et al.* 2021) for cDNA synthesis and designed an in-house real-time PCR mastermix using EvaGreen or SYBR Green I dye and in-house Hot start Taq DNA polymerase. The Ct values, amplification plots, dissociation curves were comparable between commercially available SYBR Green I PCR mastermix, and our in-house real-time PCR mastermix with either SYBR Green I or Evagreen dye. The in-house PCR mastermix is both sensitive, cost-effective and can be easily assembled. Our results therefore provide an effective solution towards developing an in-house real-time PCR mastermix which can be used as a cost-effective and efficient alternative towards commercially available real-time PCR mastermixes.

## Results

### Optimization of in-house SYBR Green I and EvaGreen PCR mastermix

Real-time PCR technique is an integration of 3 processes. RNA isolation, cDNA synthesis and real-time PCR (Fig. 1A). One of the essential steps after RNA isolation is cDNA synthesis. There are a large number of commercially available kits for cDNA synthesis which are very expensive. Here, we have used a protocol for synthesis and purification of Moloney Murine Leukemia Virus Reverse Transcriptase (M-MLV RT) (Graham *et al.* 2021) which was used for the synthesis of cDNA. Briefly, expression plasmid pET-28a_6H-MMLV_RT_D524N-6H (Addgene plasmid # 166945) was transformed into BL21 competent cells. Cultures were grown overnight at 37°C, induced with 1 mM IPTG, pelleted down and flash-frozen in liquid nitrogen and stored at −80°C until further use. Pellets were resuspended in lysis buffer and subjected for Ni-NTA purification followed by HiTrap SP sepharose purification (Fig. 1B & C). Proteins fractions were eluted in storage buffer and stored at −80°C until further use. (For buffer compositions refer materials and methods, and for detailed purification protocol refer (Graham *et al.* 2021)). cDNA synthesis was carried using 1μg of RNA from mouse ESCs as described in materials and methods. The efficiency of cDNA synthesis was analysed by GAPDH PCR (Supp. Fig. 1A). Next, cDNA was diluted to 1:10 v/v and used as template for real-time PCR. Hot-start Taq polymerase was purified as per the previously published protocol (Graham *et al.* 2021). Briefly, expression plasmid pET-28a_6H-TAQ_E602D (Addgene #166944) was transformed into BL21 competent cells, cultures were grown overnight at 37°C, induced with 1 mM IPTG, pelleted down and flash-frozen in liquid nitrogen and stored at −80°C until further use. Pellets were resuspended in lysis buffer and subjected to Ni-NTA based purification followed by HiTrap heparin purification.

**Figure 1:**
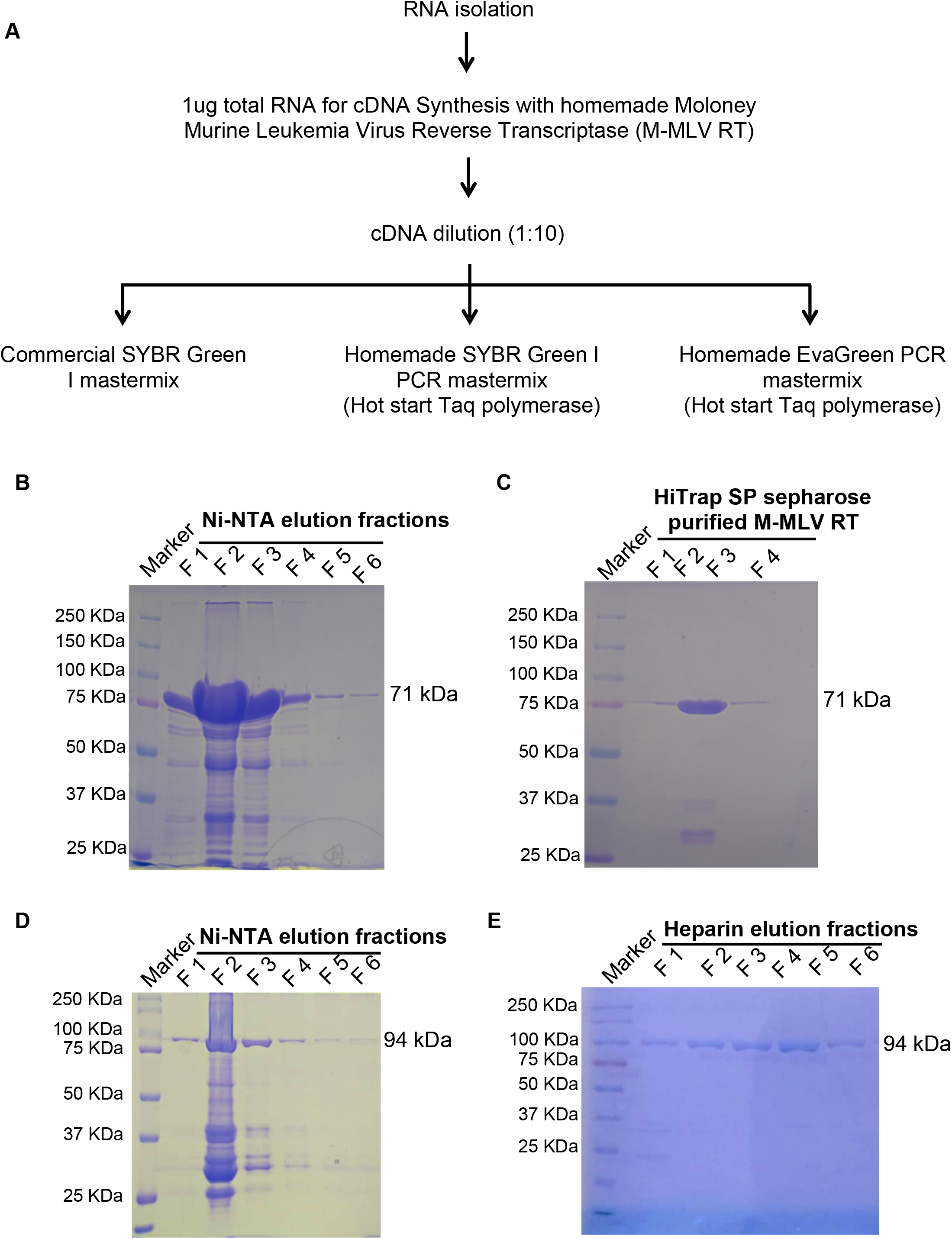
Purification of Moloney Murine Leukemia Virus Reverse Transcriptase (M-MLV RT) and Hot-start Taq polymerase: **A)** Flow chart showing various steps from RNA isolation, cDNA synthesis and real-time PCR set up under different conditions. **B)** Representative image of SDS-PAGE stained with Coomassie brilliant blue showing purified fractions F1-F6 (Fraction 1-6) after Ni-NTA purification of Moloney Murine Leukemia Virus Reverse Transcriptase (M-MLV RT). **C)** Representative image of SDS-PAGE stained with Coomassie brilliant blue showing purified fractions 1-4 of Moloney Murine Leukemia Virus Reverse Transcriptase (M-MLV RT) after HiTrap SP sepharose purification images for purification. **D)** Representative image of SDS-PAGE stained with Coomassie brilliant blue showing purified fractions F1-F6 (Fraction 1-6) after Ni-NTA purification of Hot start Taq DNA polymerase. **E)** Representative image of SDS-PAGE stained with Coomassie brilliant blue showing purified fractions F1-F6 (Fraction 1-6) after HiTrap heparin purification of Hot start Taq DNA polymerase.

Proteins fractions were eluted in storage buffer and stored at −80°C until further use (for buffer composition refer materials and methods and for detailed purification protocol refer (Graham *et al.* 2021)). This hot-start Taq polymerase was used in the preparation of the in-house PCR mastermix (Fig. 1D & E). The dNTP concentration was optimized using a final concentration of 400μM, 300μM and 250μM with EvaGreen I in-house mastermix (Supp. Table 1). Amplification was observed with all the concentrations, however there was a lot of background noise with 400 μM dNTP which was absent with 250μM dNTP (data not shown). We therefore recommend using a final concentration of 250μM dNTP for all experiments. Ct values for *Oct4*, *Sox2*, *Nestin*, *Brachyury*, *Gapdh* and *Rpl7* were consistent at all the above dNTP concentrations. Also, Ct values for duplicates of each sample were very close to each other implying the efficiency of in-house mastermixes (Supp. Table 1). For optimized brew buffer composition refer materials and methods.

### Efficiency of in-house SYBR Green I and EvaGreen PCR mastermix is comparable to commercial SYBR Green I mastermix

*Oct4, Sox2* and *Nanog* transcription factors are highly expressed in embryonic stem cells (ESCs) (Boyer *et al.* 2005). Hence, we selected transcription factors *Oct4*, *Sox2*, *Nanog* genes for real-time PCR optimization in mouse ESCs (mESCs) with *Gapdh* and *Rpl7* as housekeeping controls. Real-time PCR was set up using commercially available SYBR Green I mastermix PowerUp SYBR Green Master Mix (Applied Biosystems Cat no. A25742) and in-house SYBR Green I or EvaGreen mastermix. Ct values for *Oct4*, *Sox2*, *Nanog*, *Gapdh and Rpl7* were comparable in commercial SYBR Green I mastermix and in-house SYBR Green I or EvaGreen mastermix (Supp. Table 2, 3 & 4). Amplification plots represent the accumulation of the PCR product over successive PCR cycles. Amplification plots were comparable between the commercial SYBR Green I mastermix (Fig. 2A), and in-house SYBR Green I (Fig. 3A), or EvaGreen mastermix (Fig. 4A). Melting curve is used to analyze if there is nonspecific amplification in the PCR along with the specific band (Ririe *et al.* 1997). The single amplified product is detected by the presence of single peak whereas non-specific amplification results in appearance of multiple peaks. Melting curve for all the samples amplified with commercial SYBR Green I mastermix (Fig. 2B), and in-house SYBR Green I (Fig. 3B) or EvaGreen mastermix (Fig. 4B) had a single peak implying the presence of a single and specific amplicon in the PCR reaction.

**Figure 2:**
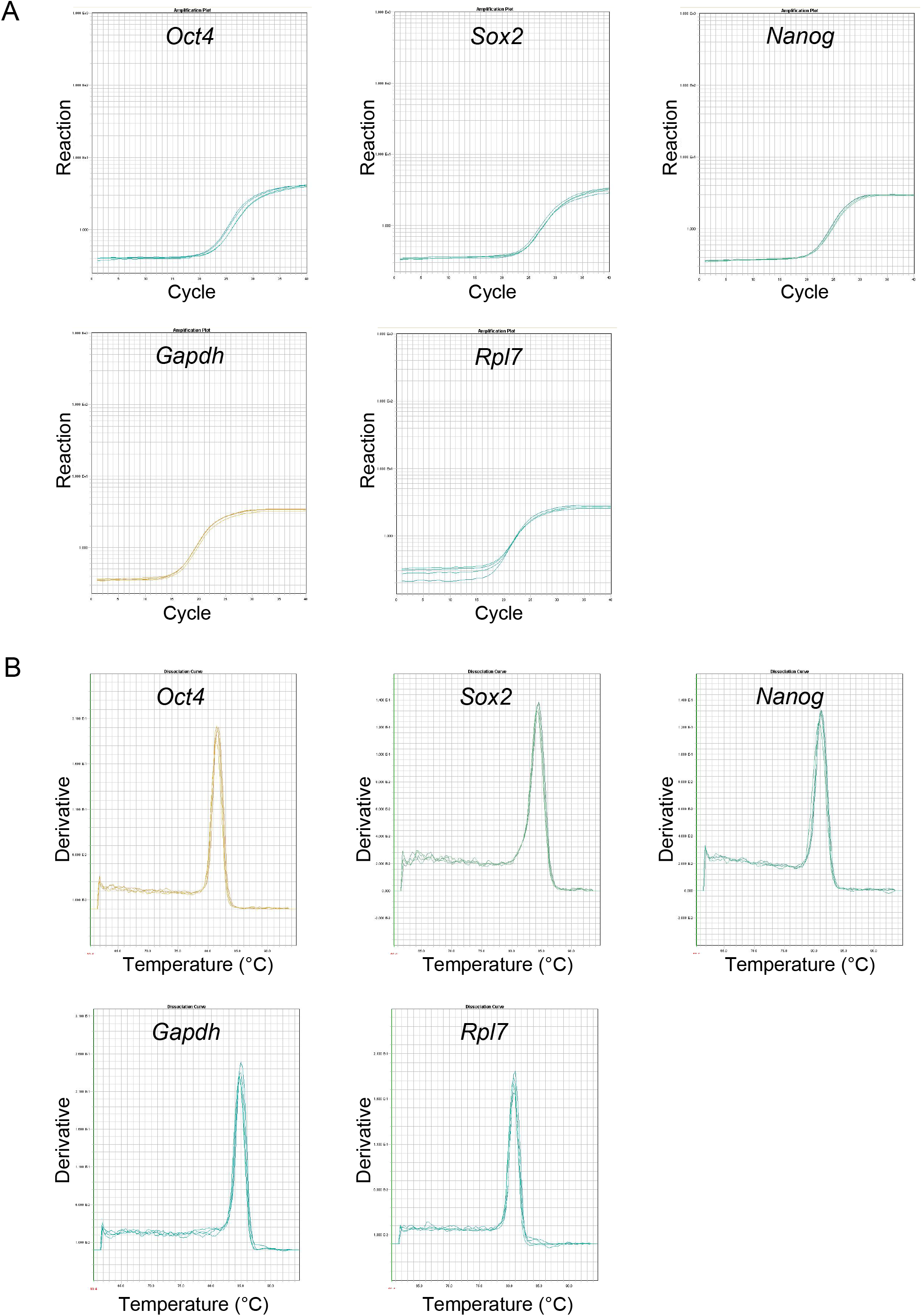
Amplification plots and dissociation curves for indicated genes with commercial SYBR Green I mastermix: **A)** Amplification plots for pluripotency marker genes *Oct4*, *Sox2*, *Nanog* and housekeeping genes *Gapdh* and *Rpl7*. **B)** Dissociation curves for pluripotency marker genes *Oct4*, *Sox2*, *Nanog* and housekeeping genes *Gapdh* and *Rpl7*.

**Figure 3:**
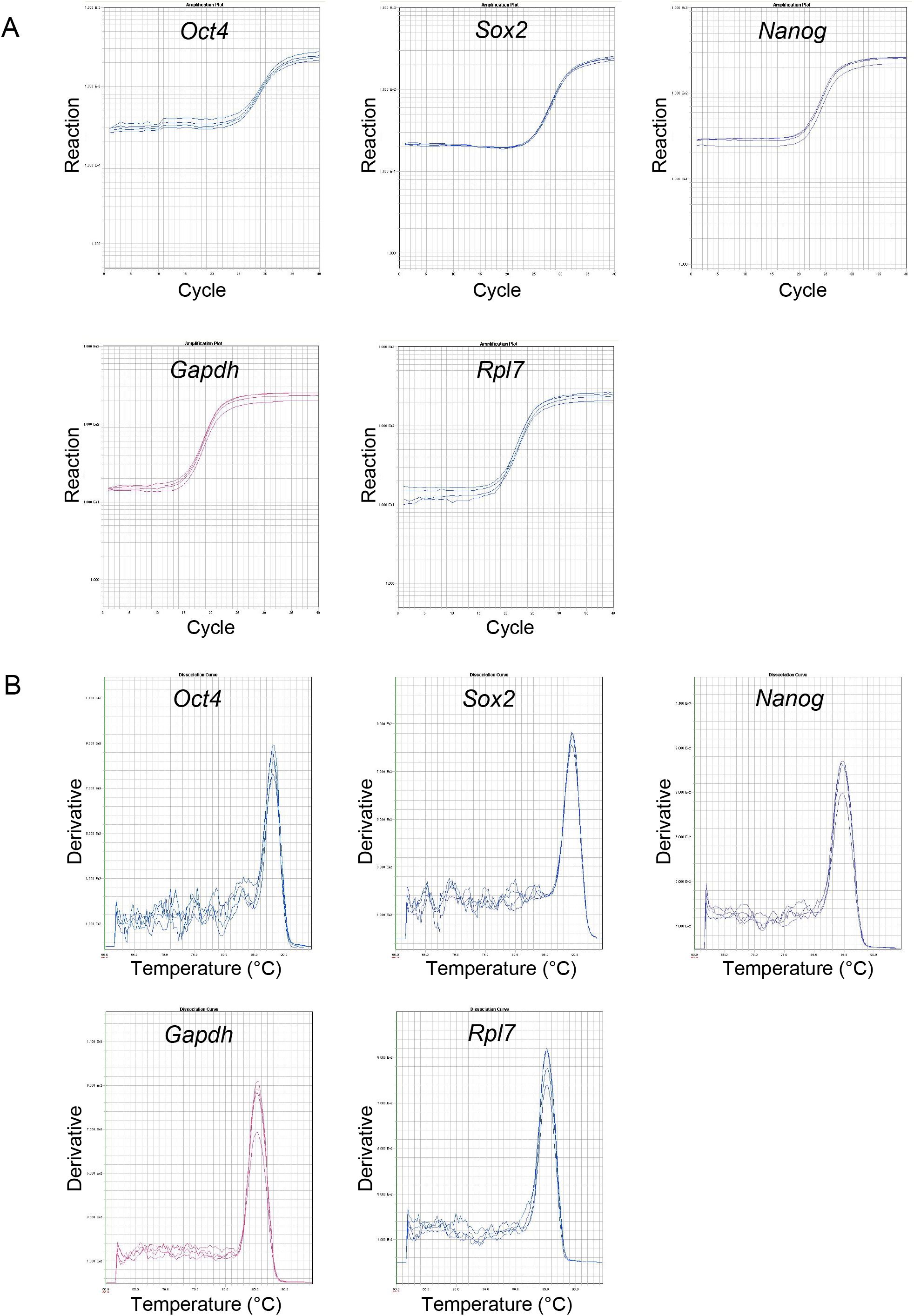
Amplification plots and dissociation curves for different genes with in-house SYBR Green I mastermix: **A)** Amplification plots for pluripotency marker genes *Oct4*, *Sox2*, *Nanog* and housekeeping genes *Gapdh* and *Rpl7*. **B)** Dissociation curves for pluripotency marker genes *Oct4*, *Sox2*, *Nanog* and housekeeping genes *Gapdh* and *Rpl7*.

**Figure 4:**
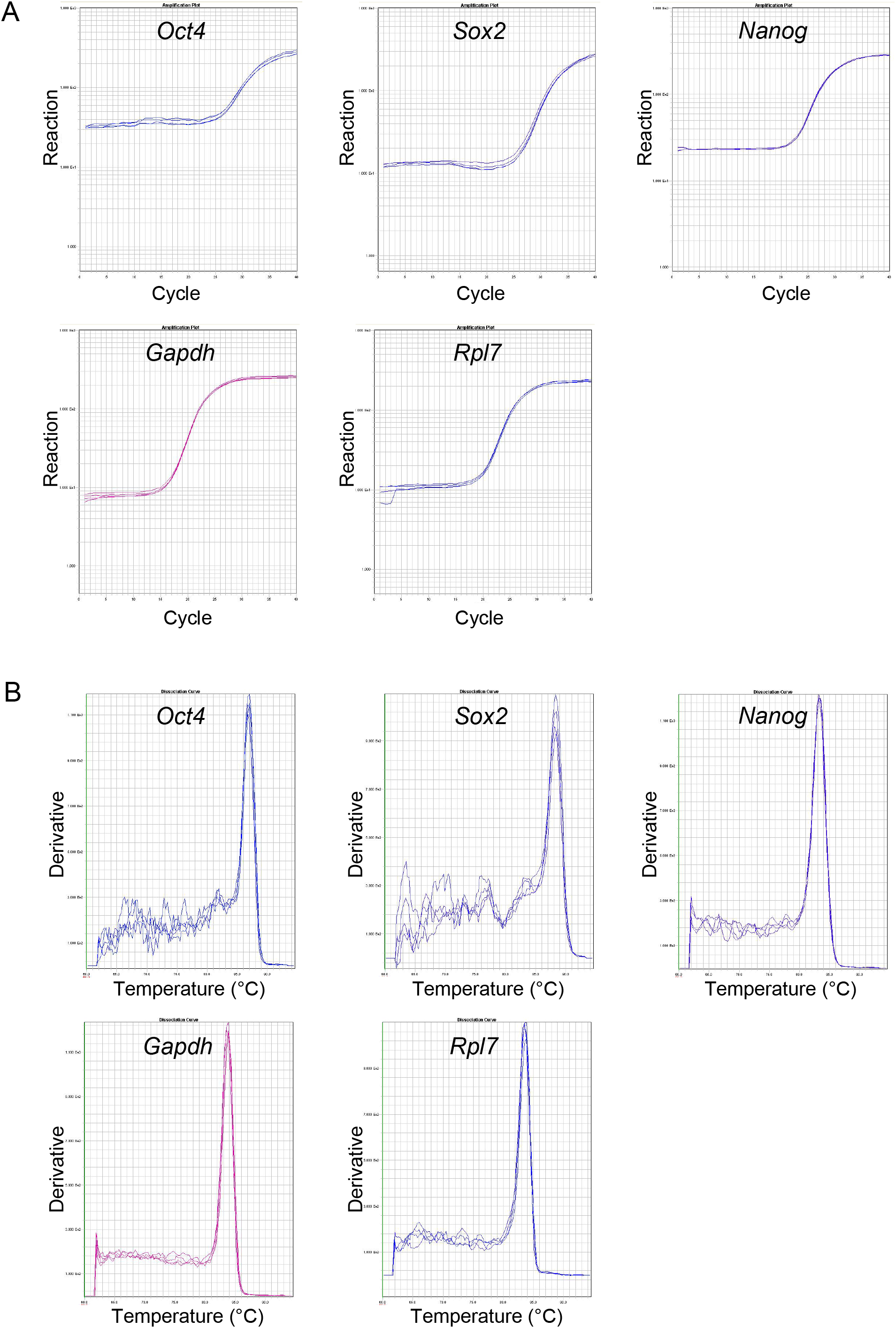
Amplification plots and dissociation curves for different genes with In-house EvaGreen PCR mastermix: **A)** Amplification plots for pluripotency marker genes *Oct4*, *Sox2*, *Nanog* and housekeeping genes *Gapdh* and *Rpl7*. **B)** Dissociation curves for pluripotency marker genes *Oct4*, *Sox2*, *Nanog* and housekeeping genes *Gapdh* and *Rpl7*.

### Costing for commercial SYBR Green I mastermix and in-house SYBR Green I or EvaGreen PCR mastermix

Real-time PCR technique consists of 3 processes. RNA isolation, cDNA synthesis and real-time PCR. RNA isolation can be done with commercially available RNA isolation kits which are very expensive. The other traditional way is to use TRIzol for RNA isolation. Details of RNA isolation using TRIzol are given in materials and methods. The next step, cDNA synthesis is done with the help of commercially available kits. A wide range of cDNA synthesis kits are available commercially with price ranging from INR 25,000 - 50,000 per 50-100 reactions. Synthesis and purification of Moloney Murine Leukemia Virus Reverse Transcriptase (M-MLV RT) for cDNA synthesis along with other components like MgCl2, dNTPs, oligo dT or random hexamers cost INR 21 per reaction. The final step is the real-time PCR set up. Commercially available SYBR Green I mastermix costs around INR 3000-4000 per ml. In-house SYBR Green I mastermix costs INR 50 per ml and EvaGreen PCR mastermix costs INR 510 per ml. For detailed costing in INR (Indian rupees) refer Supp. Table 5. In summary, we demonstrate a cost-effective and efficient method to assemble mastermixes for cDNA synthesis and RT-PCR.

## Discussion

Real-time PCR is the most favoured technique for measuring gene expression. Here, one can measure the expression of several genes with relatively low amounts of sample. However, commercially available RNA isolation kits, cDNA synthesis kits and SYBR Green I mastermix required to set up real-time PCR are very expensive and cannot be afforded by several educational institutions and small enterprises. The ongoing coronavirus pandemic has negatively impacted the research funding in a large number of developing countries, including India. This adds to the additional hurdle which limits research scholars from using expensive cutting edge techniques such as real-time PCR. Earlier, there have been attempts to prepare homemade SYBR Green I mastermix for real-time PCR. However, this mastermix contained commercial Taq polymerase and cDNA was also synthesised using commercially available kits (Karsai *et al.* 2002). This increases the expenses of conducting real-time PCR. While Graham et al demonstrate the assembly of home-made mastermixes for cDNA synthesis and real-time PCR, they do not provide a direct comparison with commercially available reagents for the same (Graham *et al.* 2021). Our study provides such a direct comparison, and also includes alternatives to SYBR Green I, such as EvaGreen. Through our manuscript, we have systematically integrated and validated protocols for RNA isolation, cDNA synthesis and real-time PCR using in-house reagents. We have compared the efficacy of these reagents with commercially available kits. Our results demonstrate that real-time PCR set up using in-house SYBR Green I or EvaGreen PCR mastermix generated Ct values, amplification plots and dissociation curves comparable to commercially available SYBR Green I PCR mastermix. These in-house reagents are not only cost-effective, but are also sensitive. Hence, these in-house reagents for real-time PCR are a promising alternative to commercially available kits.

## Materials and Methods

### RNA isolation

Total RNA was isolated from mESCs using TRIzol (Invitrogen Cat no. 15596018). Culture media was removed and cells were washed once with 1XDPBS. 500μl TRIzol was added to lyse the cells and plates were kept on the rocker for 5 minutes. Cells were scraped and the lysate was collected in an eppendorf. 100μl of chloroform was added to the TRIzol lysate and mixed thoroughly by shaking. Samples were kept at room temperature for 5 minutes and then centrifuged for 15 minutes at 12000 × g at 4°C. The upper aqueous phase containing RNA was collected in a fresh eppendorf tube and 250μl isopropanol was added to the sample. Sample was mixed and kept at room temperature for 10 minutes and later centrifuged for 10 minutes at 12000 × g at 4°C. Supernatant was discarded and the pellet was washed with 70% ethanol. Sample was centrifuged for 5 minutes at 7500 × g at 4°C. The pellet was air dried and RNase free water or DEPC-treated water was added to the sample. RNA was quantified using Nanodrop spectrophotometer.

### DNase treatment

1ug of total RNA was used for DNAseI treatment (Invitrogen cat no. 18068-015). Details of the reaction are given below.

RNA sample - 1μg

10x DNase I buffer - 1μl

DNase I - 1μl

DEPC-treated water - upto 10μl

Samples were incubated at room temperature for 15 min. 1μl of 25mM EDTA solution was added to the reaction to deactivate DNase I. Samples were kept at 65°C for 10 min followed by cDNA synthesis.

### Complementary DNA (cDNA) synthesis

DNAseI treated total RNA from mESCs was used to synthesize complementary DNA (cDNA) using random primers (Promega Cat no. C1181), dNTP mix (LAROVA Cat no. DMIX10_100ML) and in-house generated Moloney Murine Leukemia Virus Reverse Transcriptase (M-MLV RT). Details of the reaction assembly are given below.

#### cDNA synthesis reaction

5x BEAR Buffer - 4μl

10 mM dNTP - 2μl

Random Hexamers (100ng/μl) - 1μl

M-MLV Enzyme - 1μl

Dnase treated RNA reaction volume- 11 μl

Total reaction volume - 20μl

The above reaction was assembled in PCR tubes and was placed in Eppendorf mastercycler PCR machine using the following program.

PCR program:

52°C - 1 hour
95°C - 2 min

### Real time PCR

Complementary DNA (cDNA) was diluted (1:10) times and used as a template for Real-time PCR. ABI Power SYBR Green PCR master mix was used as commercial reagent. 2x in-house SYBR Green I and 2x in-house EvaGreen mix was prepared as described below. SYBR Green I (Lonza Cat no. 50513) or EvaGreen dye (Biotium Cat no. 31000), dNTP mix (LAROVA Cat no. DMIX10_100ML) were used to prepare mastermix. ABI 384 well plate (cat no. AB 1384) was used to set up real time PCR. 5μl of mastermix was added to each well. 1μl of (1:10) diluted cDNA was added as template. Total reaction volume was 6μl. ABI 7900 HT machine was used to perform Real-time PCR. Detailed composition of SYBR Green I and EvaGreen PCR mastermix is given below.

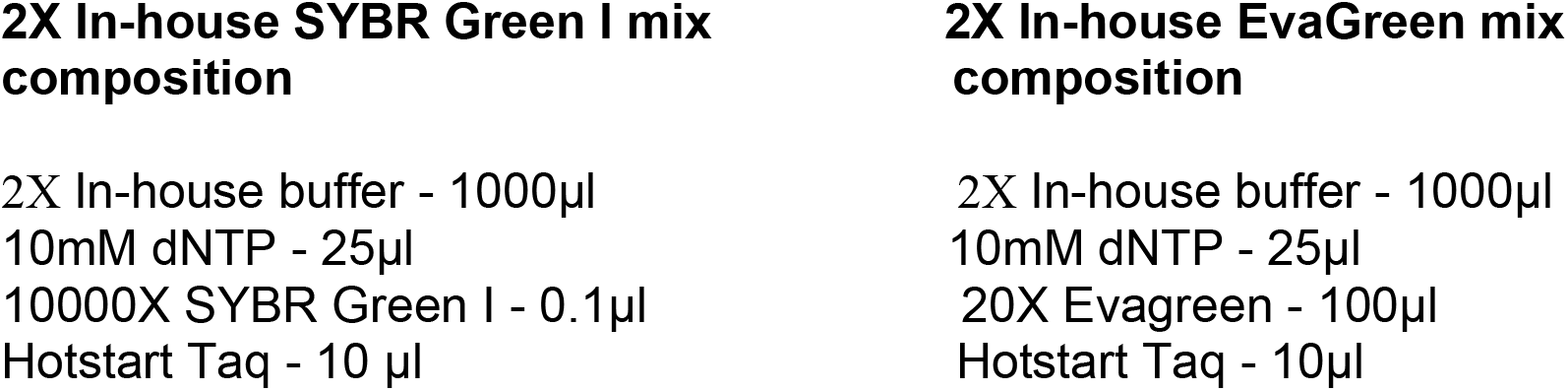

Both, 2X In-house SYBR Green I mix and 2X In-house EvaGreen mix can be stored at 4°C for up to 2 weeks. For long term storage, mastermix was stored at −20°C.

#### Real-time PCR reaction

2X In-house SYBR Green I mix or 2X In-house EvaGreen mix - 3μl

100μM Forward primer - 0.6μl

100μM Reverse primer - 0.6μl

Milli Q water - 1.88μl

Total reaction volume - 6μl

#### Real-time PCR program

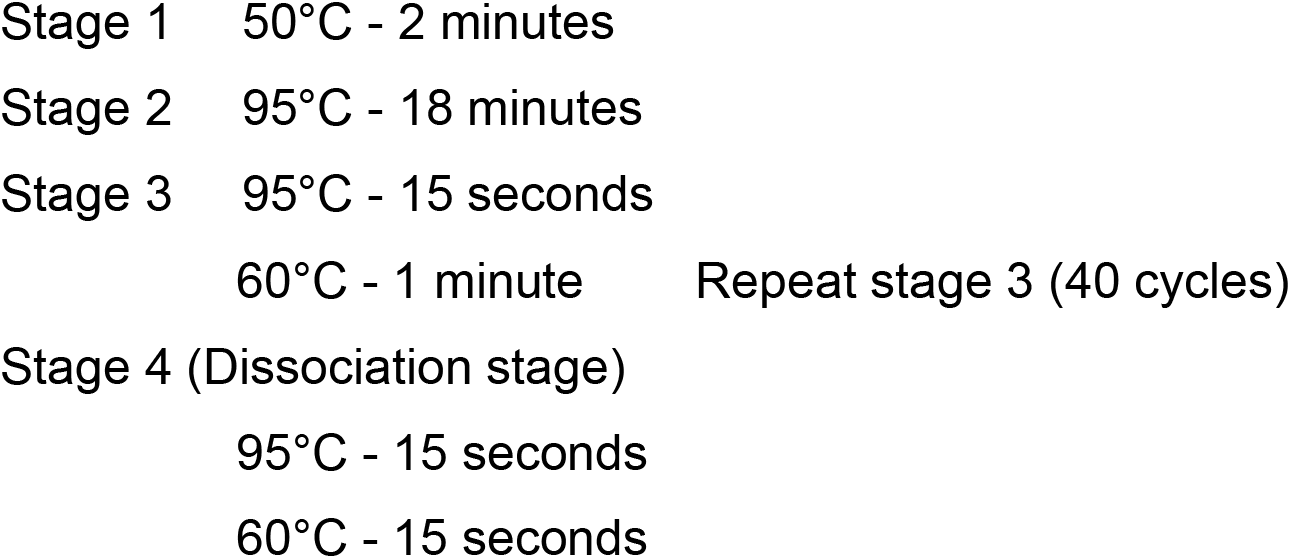

#### Buffer compositions

##### 2x In-house buffer composition-14ml

90ul 2M Tris (pH 8.1)

250ul 3M KCl

75ul 1M MgCl2

850mg Trehalose

30ul 100% Tween-20

150ul 20mg/ml BSA

##### 5x BEAR (Basic Economical Amplification Reaction) buffer composition

250 mM Tris-HCL (pH 8.4)

375 mM KCl

15 mM MgCl_2_

10% Trehalose

50 mM Dithiothreitol (DTT)

0.5mM EDTA

##### Buffers for M-MLV reverse transcriptase purification

Lysis buffer

50 mM Tris-Hcl, pH 8

100 mM NaCl

10 mM Imidazole

1 mM DTT

0.1% Triton X-100

SP buffer A

50 mM Tris-HCl, pH 8

100 mM NaCl

0.1 mM EDTA

5 mM DTT

0.1% Triton X-100

Elution Buffer

50 mM Tris-HCl, pH 8

100 mM NaCl

250 mM Imidazole

1 mM DTT

0.1% Triton X-100

M-MLV storage buffer

50 mM Tris-HCl, pH 8

100 mM NaCl

0.1 mM EDTA

5 mM DTT

0.1% Triton X-100

50% glycerol

##### Buffers for hot-start Taq polymerase preparation

Lysis buffer

50 mM Tris-HCl, pH 8

500 mM NaCl

0.1% NP-40

0.1% Triton X-100

Heparin dialysis buffer

50 mM Tris-HCl, pH 8

100 mM NaCl

0.05% NP-40

10% glycerol

5mM BME

1mM benzamidine

Taq storage buffer

50 mM Tris-HCl, pH 8

100 mM NaCl

0.1 mM EDTA 50% glycerol

3 mM DTT

---

**Supplemental Figure 1:**
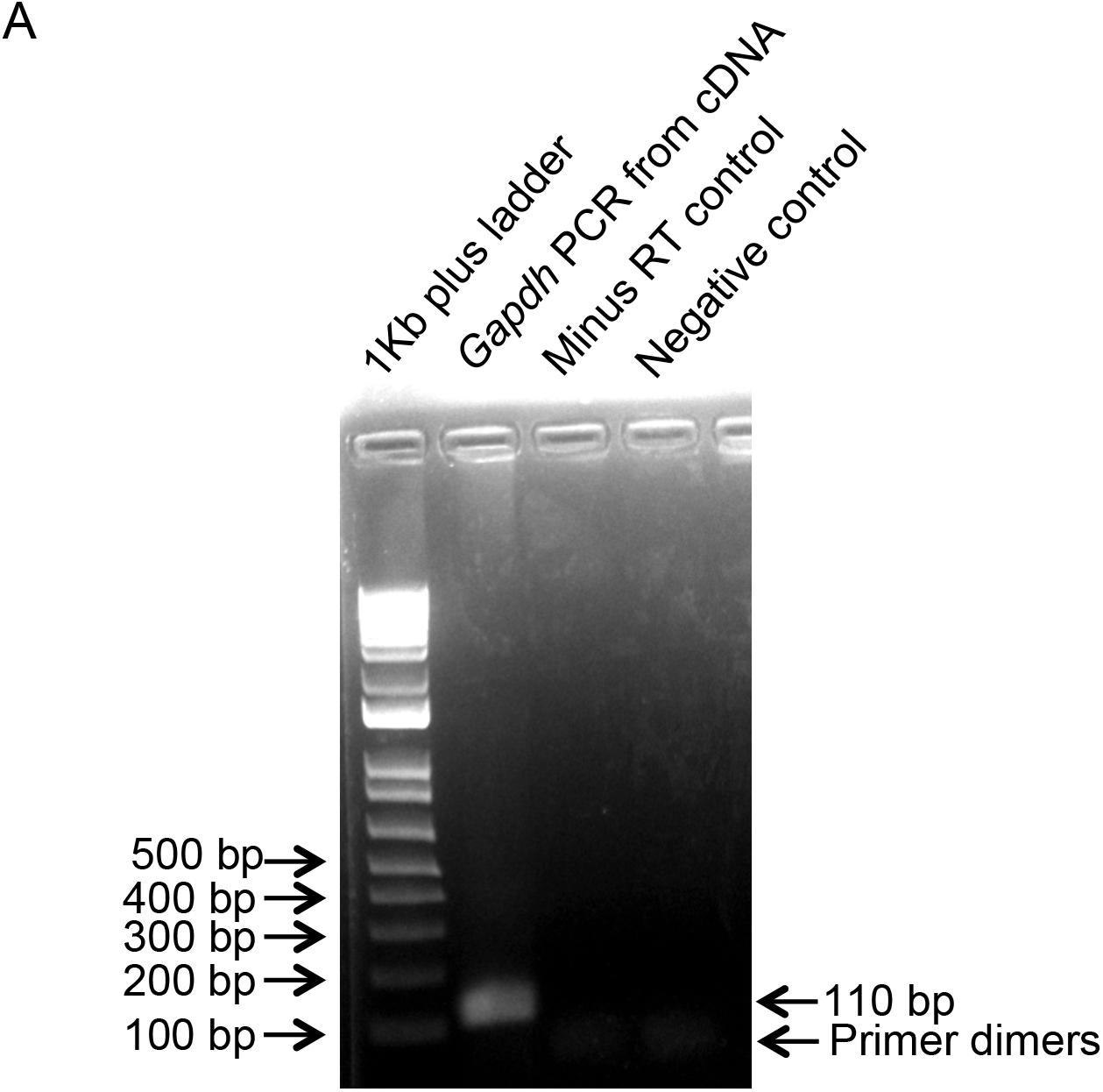
Efficiency of M-MLV reverse transcriptase **A)***Gapdh* PCR performed using mESC cDNA run on 1.5% agarose gel showing 110bp band. Marker (Lane 1), PCR using template cDNA prepared with in-house M-MLV reverse transcriptase (Lane 2), no reverse transcriptase control (Lane 3), and negative control (Lane 4).

**Supplemental Table 1:** Ct values for pluripotency marker genes *Oct4*, *Sox2*, *Nestin*, *Brachyury* and housekeeping genes *Gapdh* and *Rpl7* in mESCs with different concentrations of dNTPs 250μM, 300μM and 400μM using in-house EvaGreen PCR mastermix. S1 - Sample 1 and S2 - Sample 2. Pipetting duplicates for each sample are shown.

| Sr. No. | Sam-ple | Gene | dNTP Concen-tr-ation | Ct value | dNTP Concentr-ation | Ct value | dNTP Concentr-ation | Ct value |
| --- | --- | --- | --- | --- | --- | --- | --- | --- |
| 1 | S1 | *Oct4* | 250µM | 26.14955 | 300µM | 25.995 | 400µM | 25.98587 |
| 2 | S1 | *Oct4* | 250µM | 26.0003 | 300µM | 25.96908 | 400µM | 25.97658 |
| 3 | S2 | *Oct4* | 250µM | 26.54336 | 300µM | 26.20891 | 400µM | 26.21109 |
| 4 | S2 | *Oct4* | 250µM | 26.25872 | 300µM | 26.19582 | 400µM | 26.30681 |
| 5 | S1 | *Sox2* | 250µM | 20.50946 | 300µM | 20.9804 | 400µM | 20.79778 |
| 6 | S1 | *Sox2* | 250µM | 20.88911 | 300µM | 21.26018 | 400µM | 20.23467 |
| 7 | S2 | *Sox2* | 250µM | 21.65004 | 300µM | 21.56956 | 400µM | 21.34623 |
| 8 | S2 | *Sox2* | 250µM | 21.42214 | 300µM | 21.44382 | 400µM | 20.37359 |
| 9 | S1 | *Nestin* | 250µM | 26.9051 | 300µM | 26.82577 | 400µM | 26.78634 |
| 10 | S1 | *Nestin* | 250µM | 26.9384 | 300µM | 26.8454 | 400µM | 26.77683 |
| 11 | S2 | *Nestin* | 250µM | 27.79965 | 300µM | 27.69832 | 400µM | 27.7221 |
| 12 | S2 | *Nestin* | 250µM | 27.84153 | 300µM | 27.76137 | 400µM | 27.67209 |
| 13 | S1 | *Brachyury* | 250µM | 25.02074 | 300µM | 25.25898 | 400µM | 25.1886 |
| 14 | S1 | *Brachyury* | 250µM | 24.86784 | 300µM | 24.82618 | 400µM | 25.08578 |
| 15 | S2 | *Brachyury* | 250µM | 27.85045 | 300µM | 26.53506 | 400µM | 27.22934 |
| 16 | S2 | *Brachyury* | 250µM | 26.57951 | 300µM | 26.58253 | 400µM | 27.76588 |
| 17 | S1 | *Gapdh* | 250µM | 18.59947 | 300µM | 18.50684 | 400µM | 18.35302 |
| 18 | S1 | *Gapdh* | 250µM | 18.64362 | 300µM | 18.95817 | 400µM | 18.41865 |
| 19 | S2 | *Gapdh* | 250µM | 19.30496 | 300µM | 19.1272 | 400µM | 19.05853 |
| 20 | S2 | *Gapdh* | 250µM | 19.23951 | 300µM | 19.12005 | 400µM | 19.07347 |
| 21 | S1 | *Rpl7* | 250µM | 21.03698 | 300µM | 20.92539 | 400µM | 20.84437 |
| 22 | S1 | *Rpl7* | 250µM | 21.08816 | 300µM | 20.95721 | 400µM | 20.81063 |
| 23 | S2 | *Rpl7* | 250µM | 21.89484 | 300µM | 21.72218 | 400µM | 21.58777 |
| 24 | S2 | *Rpl7* | 250µM | 21.82392 | 300µM | 21.71718 | 400µM | 21.58092 |

**Supplemental Table 2:** Ct values for pluripotency marker genes *Oct4*, *Sox2*, *Nanog* and housekeeping genes *Gapdh* and *Rpl7* in mESCs with commercial SYBR Green I mastermix.

| Sr. No. | Sample | Gene | Dye | Ct value |
| --- | --- | --- | --- | --- |
| 1 | mESCs Sample 1 | *Oct4* | ABI SYBR | 20.68661 |
| 2 | mESCs Sample 2 | *Oct4* | ABI SYBR | 21.54116 |
| 3 | mESCs Sample 1 | *Sox2* | ABI SYBR | 22.94617 |
| 4 | mESCs Sample 2 | *Sox2* | ABI SYBR | 23.39544 |
| 5 | mESCs Sample 1 | *Nanog* | ABI SYBR | 20.02914 |
| 6 | mESCs Sample 2 | *Nanog* | ABI SYBR | 20.40491 |
| 7 | mESCs Sample 1 | *Gapdh* | ABI SYBR | 15.0164 |
| 8 | mESCs Sample 2 | *Gapdh* | ABI SYBR | 15.25804 |
| 9 | mESCs Sample 1 | *Rpl7* | ABI SYBR | 17.7328 |
| 10 | mESCs Sample 2 | *Rpl7* | ABI SYBR | 17.96429 |

**Supplemental Table 3:** Ct values for pluripotency marker genes *Oct4*, *Sox2*, *Nanog* and housekeeping genes *Gapdh* and *Rpl7* in mESCs with in-house SYBR Green I mastermix.

| Sr. No. | Sample | Gene | Dye | Ct value |
| --- | --- | --- | --- | --- |
| 1 | mESCs Sample 1 | Oct4 | SYBR Green I | 18.94675 |
| 2 | mESCs Sample 2 | Oct4 | SYBR Green I | 19.37616 |
| 3 | mESCs Sample 1 | Sox2 | SYBR Green I | 19.85928 |
| 4 | mESCs Sample 2 | Sox2 | SYBR Green I | 18.6483 |
| 5 | mESCs Sample 1 | Nanog | SYBR Green I | 13.19007 |
| 6 | mESCs Sample 2 | Nanog | SYBR Green I | 12.85204 |
| 7 | mESCs Sample 1 | Gapdh | SYBR Green I | 11.24917 |
| 8 | mESCs Sample 2 | Gapdh | SYBR Green I | 12.45861 |
| 9 | mESCs Sample 1 | Rpl7 | SYBR Green I | 12.18782 |
| 10 | mESCs Sample 2 | Rpl7 | SYBR Green I | 12.99236 |

**Supplemental Table 4:** Ct values for pluripotency marker genes *Oct4*, *Sox2*, *Nanog* and housekeeping genes *Gapdh* and *Rpl7* in mESCs with in-house EvaGreen PCR mastermix.

| Sr. No. | Sample | Gene | Dye | Ct value |
| --- | --- | --- | --- | --- |
| 1 | mESCs Sample 1 | *Oct4* | Evagreen | 20.91577 |
| 2 | mESCs Sample 2 | *Oct4* | Evagreen | 21.43513 |
| 3 | mESCs Sample 1 | *Sox2* | Evagreen | 19.69726 |
| 4 | mESCs Sample 2 | *Sox2* | Evagreen | 20.38235 |
| 5 | mESCs Sample 1 | *Nanog* | Evagreen | 16.11342 |
| 6 | mESCs Sample 2 | *Nanog* | Evagreen | 14.94121 |
| 7 | mESCs Sample 1 | *Gapdh* | Evagreen | 12.86926 |
| 8 | mESCs Sample 2 | *Gapdh* | Evagreen | 11.67245 |
| 9 | mESCs Sample 1 | *Rpl7* | Evagreen | 14.91321 |
| 10 | mESCs Sample 2 | *Rpl7* | Evagreen | 15.093 |

**Supplemental Table 5:** Costing evaluation of RNA isolation, cDNA synthesis and real time PCR set up using commercially available kits and by in-house reagents.

| Sr. No. | Process | Commercial kit cost (in Indian rupees INR) | In-house reagent cost (in Indian rupees INR) | % Savings with in-house reagents |
| --- | --- | --- | --- | --- |
| 1 | RNA isolation | INR 300-400 per sample | INR 62.5 using TRIzol | 79.16 – 84.37% |
| 2 | cDNA synthesis | INR 260 per reaction | INR 21 per reaction<br>(Includes cost of beads, SP sepharose, Ni-NTA, plastic columns to be packed, dialysis tubing, buffers, LB broth, MgCl<sub>2</sub>, dNTPs, random hexamers or oligo dT | 91.92% |
| 3 | Real time PCR SYBR Green I mastermix | INR 3000 – 4000 per ml | INR 50 per ml for SYBR GreenI mastermix<br>INR 510 per ml for Evagreen I mastermix | 98.33 – 98.75%<br>83 – 87.25% |

## Acknowledgements

This work was supported by funds to D.S. from the Department of Biotechnology (BT/PR25883/GET/119/105/2017), and ICMR (2020-3076/SCR/ADHOC-BMS) D.S, A.M., V.S., V.T. acknowledge intramural funding from National Centre for Cell Science. S.B.S is a recipient of a Senior Research Fellowship from the Department of Biotechnology, India; M.T. and J.S. are recipients of a Senior Research Fellowship from University Grants Commission, India; H. is a recipient of a Senior Research Fellowship from CSIR, India. We thank members of the Subramanyam and Majumdar lab for constructive discussion.

## Author contributions

A.M, D.S and R.D.M. conceived and designed the study. R.D.M. contributed to experiments in Fig. 1A, 2, 3, 4, Fig. S1, Table S1–S4. A.M, S.B.S, H. and J.S contributed to experiments in Fig. 1B-E. M.T contributed to experiments in Fig. 3, 4. D.S and R.D.M wrote the manuscript with help from A.M, V.S., V.T and S.L.V. All authors reviewed the manuscript.

